# *In vivo* calcium imaging shows that satellite glial cells have increased activity in painful states

**DOI:** 10.1101/2022.10.07.511264

**Authors:** Sara E. Jager, George Goodwin, Kim I. Chisholm, Franziska Denk

## Abstract

Satellite glial cells (SGCs) are important for proper neuronal function of primary sensory neurons to whom they provide homeostatic support. Most research of SGC function has been performed with *in vitro* studies, but recent advances in calcium imaging and transgenic mouse models have enabled this first *in vivo* study of single cell SGC function in mouse models of inflammation and neuropathic pain.

We found that in naïve condition SGCs do not respond in a time-locked fashion to neuronal firing. In painful inflammatory and neuropathic states we detected time-locked signals in a subset of SGCs, but only with suprathreshold stimulation of the sciatic nerve. Surprisingly therefore, we conclude that most calcium signals in SGCs seem to develop at arbitrary intervals not directly linked to neuronal activity patterns.

More in line with expectations, our experiments also revealed that the number of active SGCs was increased under conditions of inflammation or nerve injury. This could reflect the increased requirement for homeostatic support across dorsal root ganglion neuron populations, which are more active during such painful states.

## Introduction

Our ability to feel touch, temperature or pain is reliant on sensory neurons which connect our spinal cord and brain to the environment. What is often overlooked, however, is that these neurons cannot function in isolation but require homeostatic support from Schwann cells and satellite glial cells (SGCs). The cell bodies of our peripheral sensory neurons are enveloped by SGCs ^1,2^; in rodents it appears that the larger the neuron, the more SGCs it has – with up to ten reported per cell ^3^.

SGCs are believed to provide buffering functions, e.g. by taking up potassium ^4,5^ and neurotransmitters from the extracellular space surrounding the neurons ^6^. SGCs are thus crucial for maintaining homeostasis, as sensory neurons require tight environmental control in order to maintain their membrane potential and firing properties. There is also evidence to suggest that SGC function changes when sensory neurons are injured or affected by disease ^7^. For example, studies report that SGCs show increased expression of gap junction transcripts and proteins after traumatic nerve injury, strengthening inter-SGC connectivity ^4,8–10^. These changes could be a reflection of SGCs responding to altered neuronal function in an injured or diseased environment.

Despite their pivotal contribution to somatosensation, many crucial questions about SGC communication remain unanswered. The primary reason for this is likely that they are very difficult to study. Without antibodies, SGCs are hardly visible, which has meant that their presence in peripheral neuronal culture systems often goes unnoticed ^11^. On the other hand, SGCs cultured in the absence of neurons rapidly regress into a Schwann-cell like precursor phenotype ^12,13^, which means that any *in vitro* studies of pure SGCs ideally have to be undertaken within a few days of extraction. Luckily, progress in optical imaging techniques has meant that SGC function can now finally be studied *in vivo*, using calcium imaging of dorsal root ganglia (DRG) ^14^, the site of peripheral neuron-SGC units.

Here, we used a transgenic approach, using *Fabp7*, a SGC marker gene ^15^, to drive expression of the calcium sensor GCaMP6s. The transgenic mouse we chose has previously been used in the SGC field, seemingly successfully^15,16^. Equally, GCaMP6s imaging has been increasingly used to assess cellular activity in DRG *in vivo* ^17–20^. Here, we set out to use these techniques to investigate the following key questions:

1. Do SGCs generate calcium transients in response to peripheral neuron stimulation? We know from *in vitro* experiments that ATP is released by neuronal cell bodies in an activity-dependent manner ^21^. The ATP has then been shown to bind to purinergic receptors on SGCs, where it results in an intracellular calcium signal which affects downstream cellular functions, including communication with neurons ^21,22^. The resulting calcium waves can be further propagated to neighboring SGCs or neurons via gap junctions ^23^. However, to date, all of this work has been conducted *in vitro*, and it is therefore not clear whether neuronal activity can also generate calcium signals in SGCs in a physiological, *in vivo* situation.
2. Do SGCs increase their responses when the peripheral nervous system is sensitized, e.g. by inflammation or injury? This has frequently been suggested based on *in vitro* studies. Thus, stimulation with ATP was shown to result in a higher calcium signal in primary SGC cultures derived from inflammatory models ^24–26^. Most of the other literature is based on indirect measures of SGC activation. For example, many have observed an increase in gap junctions between SGCs upon sensitization ^4,8–10^, which should allow for increased propagation of signals between cells. Others, including ourselves, have identified molecular changes that indicate increased SGC responses in disease states, such as alterations in mRNA expression related to cholesterol ^27^ and lipid metabolism ^15^ and regulation of various proteins such as rat GFAP ^28,29^, Hmgcs1 ^30^ and Kir4.1 ^31,32^. In this paper we conduct *in vivo* calcium imaging to obtain a direct measure of whether 1) neuronal activity can induce calcium transients in healthy SCGs and 2) whether inflammation or neuropathy can increase this signalling mechanism.

## Results

First, we validated the *Fabp7*-CreER driver line to ensure that it is specific for SGCs. For this, we generated two crosses: *Fabp7*-CreER with flox-STOP-*GCaMP6s* for immunofluorescence and calcium imaging experiments, and *Fabp7*-CreER with the nuclear-tagged reporter line flox-STOP-*Sun1GFP*. The latter reporter line is a superior tool for flow cytometry compared to the GCaMP6s reporter, since its green fluorescent protein is constitutively active. We found that *Fabp7*-CreER is only a very weak driver of recombination, with only about half of all tamoxifen-treated mice showing the expected reporter gene expression. Mice with insufficient reporter gene labelling were excluded from data collection and/or analysis. Immunostaining of DRGs from successfully recombined *Fabp7*-CreER-*GCaMP6s* mice (*n* = 6-7) showed that an average of 76.5% of GCaMP6s positive cells were also positive for the SGC marker glutamine synthetase (GS, Fig. 1A-C) and that 85% of GCaMP6s positive cells were positive for Fabp7 (Fig 1D-E). For the GS staining similar results were obtained via flow cytometry of *Fabp7*-CreER-*Sun1GFP* mice (*n* = 5), revealing 68.5% GS positivity among GFP+ cells (Fig. 1F-I, Suppl. Fig. 3A).

**Figure 1:**
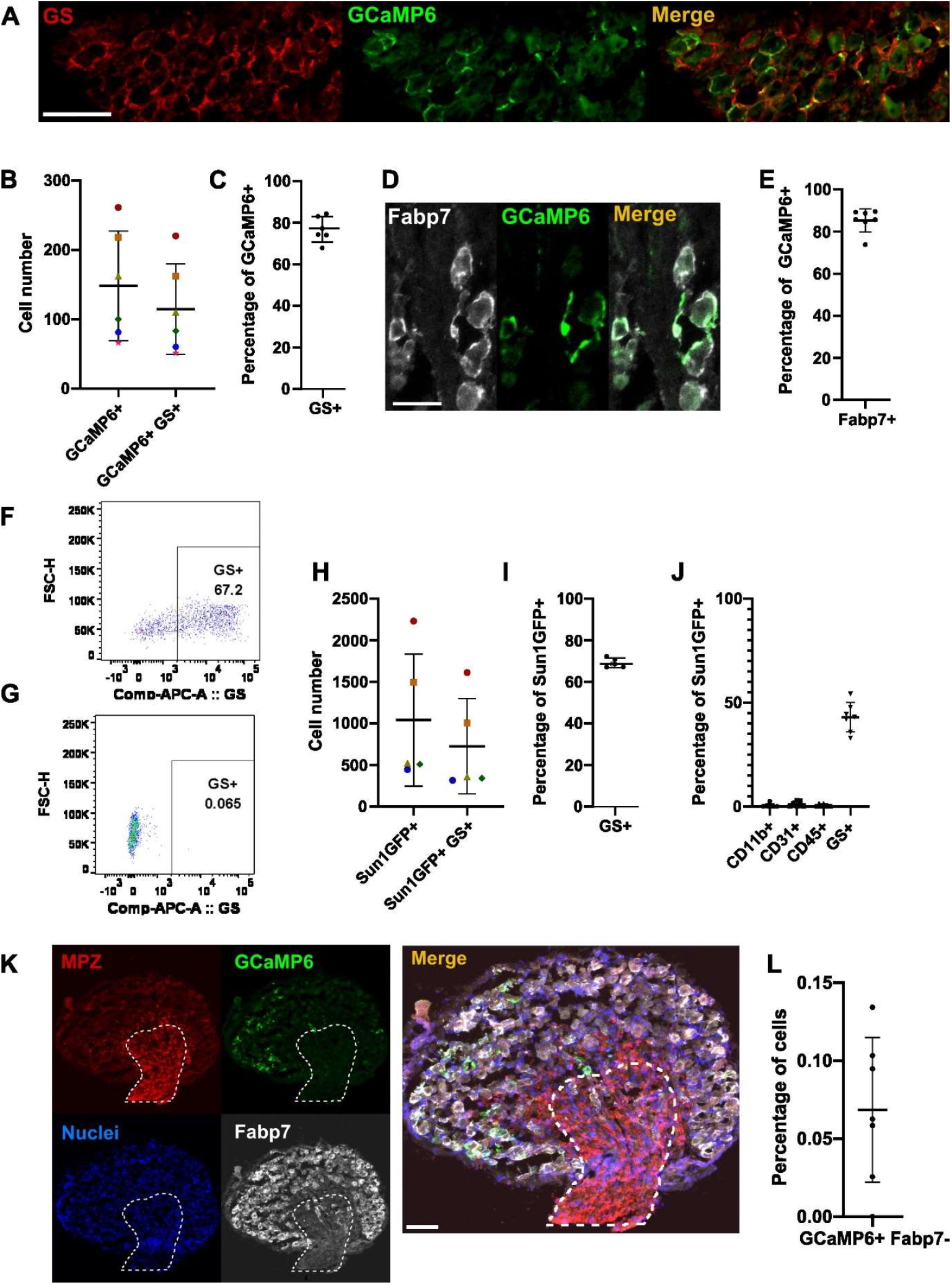
Using Fabp7-CreER-GCaMP6s mice to target a calcium sensor to SGCs. **(A)** Representative image of immunofluorescent staining of a DRG from the Fabp7-CreER-GCaMP6s line. SGCs were visualised with an antibody against the SGC marker glutamine synthetase (GS, red), while GCaMP6s signal was enhanced with an antibody against green fluorescent protein (GFP, green). Scale bar = 100 μm. **(B)** Quantification of immunofluorescence. Shown are the absolute counts of GCaMP6+ positive cells and GCAMP6+/GS+ double positive cells. Data from n = 6 mice (same colour = data from the same animal). Error bars = standard of deviation. Six sections were counted for each mouse, and the counts added to feed into this plot. Black lines indicate the mean value. **(C)** Plot of the percentage of GCaMP6s+ cells also positive for GS+, calculated from the values in (B). Error bars = standard of deviation **(D)** Representative image of immunofluorescent staining of DRG tissue from the Fabp7-CreER-GCaMP6s line. SGC visualized with an antibody against Fabp7 (grey) while GCaMP6s signal was enhanced with an antibody against GFP (green). Scale bar = 50 μm. (**E**) Quantification of percentage of GCaMP6s+ cells also positive for Fabp7. Data from n = 7 mice, 4 sections counted for each mouse. Black lines indicate the mean value and error bars = standard of deviation. (**F**) Representative image of flow cytometry result of L3-L5 DRGs dissociated from the Fabp7-CreER-Sun1GFP line, with SGCs stained for GS. The events shown in the plot are single live cells positive for GFP. See Suppl. Fig. 3A for full gating strategy. **(G)** Fluorescence-minus one control for (F), showing the same sample without GS+ antibody stain. **(H)** Quantification of absolute numbers of Sun1GFP-expressing cells measured via flow cytometry of L3-L5 DRG, n = 5 mice. Black lines indicate the mean value. Error bars = standard of deviation **(I)** Plot of the percentage of Sun1GFP+ cells also positive for GS+ (n = 5 mice). Error bars = standard of deviation. **(J)** Quantification of flow cytometric results from dissociated DRGs from the Fabp7-CreER-GCaMP6s line, stained with markers for myeloid cells (CD11b+), endothelial cells (CD31+), immune cells (CD45+) and SGCs (GS+). Black lines indicate mean value. Error bars = standard of deviation. See Suppl. Fig 4 for full gating strategy. **(K)** Representative image of immunofluorescent staining experiment designed to probe for off-target GCaMP6s signal in the nerve trunk: myelin protein zero in red (MPZ), GFP+ GCaMP6s in green, DAPI+ nuclei in blue and Fabp7+ SGCs in white. The encircled area is identified based on the high immunoreactivity for MPZ and is where the nerve enters the DRG. This region is rich in axons, fibroblasts and Schwann cells. Scale bar = 100 μm. **(L)** Quantification of the percentage of GCaMP6+/Fabp7-cells found in the area stained by MPZ. The number of cells was determined by counting the number of nuclei. Four sections were counted per mouse, and the resulting percentages plotted here from n = 7 mice. Error bars = standard of deviation.

To test for off-target expression of the *Fabp7*-CreER driver, we performed further flow cytometry and immunofluorescence experiments. With flow cytometry, amongst the GFP+ population, 0.41% of cells were also positive for the myeloid cell marker CD11b, 1.1% for the endothelial cell marker CD31 and 0.5% for the pan-immune cell marker CD45 (Fig. 1J). Using this more extensive flow panel (Suppl. Fig. 4), we detected fewer GS+ cells among the Sun1-GFP+ population (44% on average). We suspect that this was due to spectral interference which reduced our precision for distinguishing GS+ from GS-cells in this particular experimental run. Using immunostaining, we checked for GCaMP6s expression in the area where the nerves enter the DRG (Fig. 1K). On average we found 0.07% of the cells in this area to be GCaMP6+Fabp7-(n=7) (Fig. 1L), suggesting that there is little or no off-target expression in Schwann cells and fibroblasts. The specificity of the expression is in line with data previously presented by Avraham et al. and Mapps et al. ^15,16^.

Overall, these analyses indicated that any GCaMP6s+/GS-cells are likely to represent GS negative SGCs ^15^, rather than ectopic expression of GCaMP6s in other cell types. However, while we can thus assume with reasonable confidence that GCaMP6+ cells are SGCs in *Fabp7*-CreER-*GCaMP6s* DRG, the same cannot be said for the reverse: in our hands many SGCs in *Fabp7*-CreER mice were not positive for GCaMP6s and/or Sun1GFP. Specifically, in the flow cytometric data on average only 7.8% of GS positive SGCs were also positive for Sun1GFP (Suppl. Fig. 3B). For both lines we found very sparse and variable labelling, likely due to the lack of the T2 mutations that were introduced into the CreER fusion protein by Pierre Chambon and Daniel Metzger in the late 1990s ^40,41^. Bearing these restrictions in mind, we proceeded to *in vivo* calcium imaging experiments. Calcium imaging of lumbar L4 DRG was performed in live anaesthetised mice using an upright confocal/multiphoton microscope with a fully open pinhole, as described previously (Fig. 2A) ^20^. In well-labelled mice (24 mice out of 60), GCaMP6s+ cells could readily be identified by eye and had clear SGCs morphology (Fig. 2B). Calcium signal was readily discernible against background in most cells (Fig. 2C and Suppl. videos), with only 13 out of 1418 SGCs displaying baseline activity high enough to prevent the use of our set response threshold. The threshold was determined based on visual inspection of our videos to ensure that only true fluorescence signal would be included in subsequent analyses. It was set as signal at 25%+2*standard deviations above the mean cell fluorescence over the entire recording period or until injection of pentobarbital. Plotting responses across *n* = 6 naïve animals (2 males, 4 females), we could not detect any discernible connection between SGC calcium signal and neuronal stimulation (Fig. 2D and Suppl. Fig. 1). Instead, it seemed that they responded at arbitrary intervals throughout the recording period regardless of neuronal activity.

**Figure 2:**
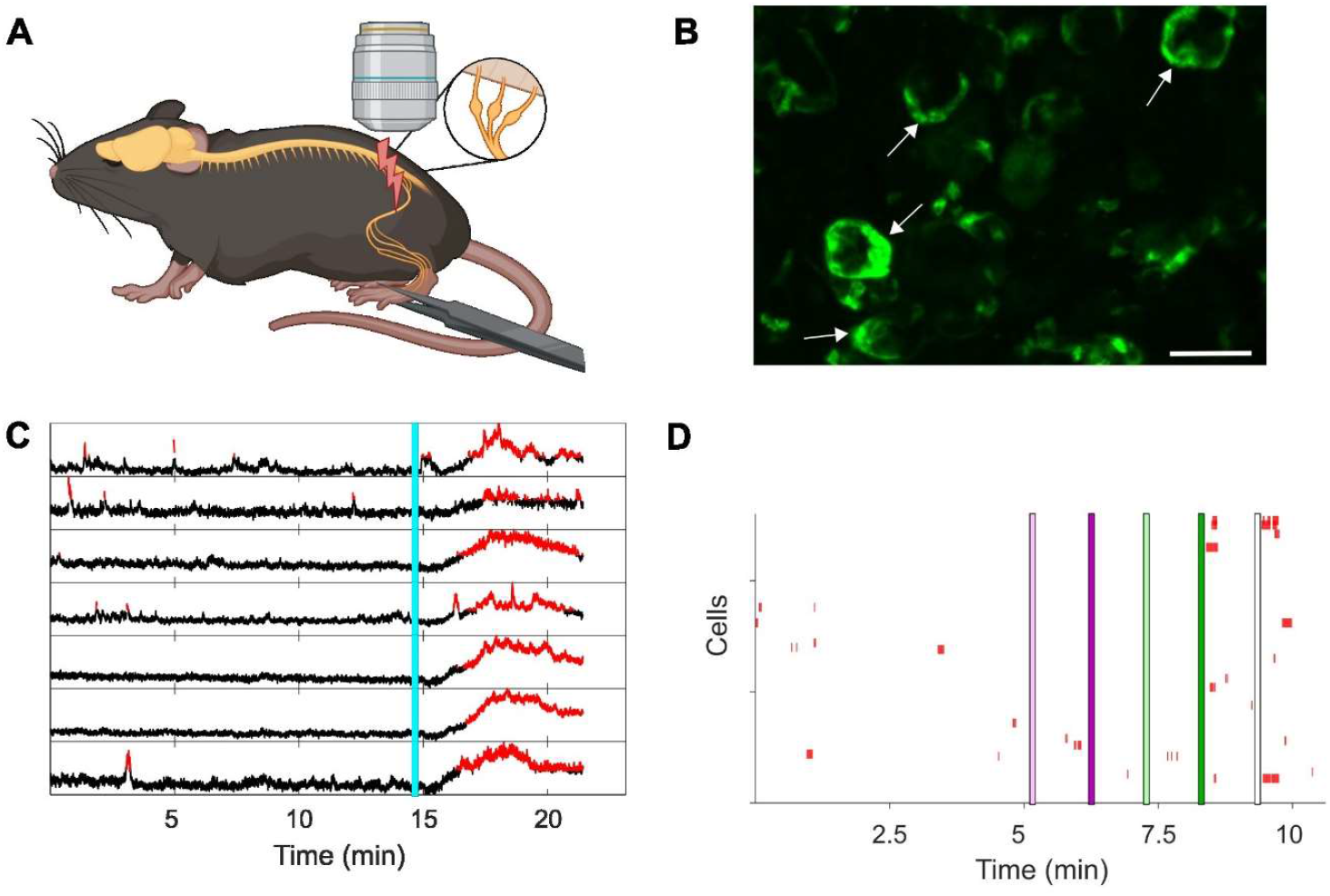
In vivo Ca^2+^ imaging of SGCs. **(A)** Diagram of the experimental setup. Fabp7-CreER-GCaMP6s mice were anaesthetised, their L4 DRG exposed and imaged using an upright confocal/multiphoton microscope as described previously^20^. The enlargement shows L3-L5 DRGs that contain the cell bodies of neurons within the sciatic nerve. Neurons were activated using electrical and natural stimuli. Created with BioRender.com. **(B)** View through the microscope during imaging at 20x magnification. Scale bar = 50μm. The white arrows point to clear examples of SGCs surrounding neuronal cell bodies. **(C)** Representative fluorescent traces from seven individual SGCs during an imaging session. The turquoise line indicates injection of an overdose of pentobarbital resulting in a post-mortem influx of calcium. The red parts of the traces show when the fluorescence signals are higher than the background indicating a response. The threshold was set as 25% above the mean + 2*standard deviation based on visual inspection of the recorded videos. **(D)** Representative raster plot of all SGCs recorded from one mouse (for this mouse n=127 cells). Each red colour annotation represents a significant Ca^2+^ signal in an individual SGC. Coloured lines (pink, green or white) indicate times at which neurons were stimulated: single A fibre stimulation (light pink line), 10 sec of A fibre stimulation (5 Hz pulses, dark pink line), single suprathreshold fibre stimulation (light green line), 10 sec of suprathreshold stimulation (5 Hz pulses, dark green line), squeezing of the paw (5 sec, white line).

This changed somewhat once peripheral neurons were sensitized, either through injection of an inflammatory stimulus into the paw (Complete Freund’s Adjuvant, CFA, 20μl) or after traumatic nerve injury as a result of partial sciatic nerve ligation (pSNL). One day after CFA or seven days after pSNL, we observed a higher number of time-locked calcium responses than expected by chance, but only after repeated suprathreshold stimulation (Table 1). This was not the case for either A fibre, single suprathreshold or paw pinch stimulation.

**Table 1:** Table showing the likelihood of observing a time-locked response by chance (“Expected by chance”) and the actual observed responses (“Observed in data”) in the collected data. “Chance per cell” shows the theoretical chance that one cell appears time-locked based on the frequency of the responses and the length of the recording. “Number of detected cells” states the number of cells recorded for each condition in all videos. “Expected by chance in data” is the number of times a cell will appear time-locked by chance. “Observed in data” is the number of times a time-locked response was recorded in the data after electrical stimulation or paw squeeze.

|  | Chance per cell | Number of detected cells | Expected by chance in data | Observed in data: A fiber | Observed in data: A fiber repeated | Observed in data: Suprathreshold | Observed in data: Suprathreshold repeated | Observed in data: Paw squeeze |
| --- | --- | --- | --- | --- | --- | --- | --- | --- |
| Naïve | 0.011 | 368 | 4 | 2 | 2 | 2 | 3 | 3 |
| CFA | 0.0075 | 533 | 4 | 2 | 2 | 2 | 11 | 5 |
| pSNL | 0.014 | 517 | 7 | 4 | 5 | 4 | 9 | 5 |

Next, we looked across the entire recorded period to determine whether each SGC was, at any timepoint, responding with a calcium signal (Fig. 3A-B). Our pain models induced an increase in the percentage of responding SGCs (Fig. 3C), with the mean number of responders rising from 17% in naïve mice (*n* = 6) to 32.6% in CFA mice (d = 1.25, *n* = 6) and 40% in pSNL mice (d = 1.84, *n* = 7). This change was statistically significant for the pSNL vs. naïve comparison using an ordinary one-way ANOVA test followed by Tukey’s multiple comparisons test of means (*p* = 0.01). This change was not due to a change in the total number of SGCs that were included in the analysis of any given group (Fig. 3D). Moreover, sensitization did not appear to increase the frequency of responses in any individual SGC (Fig. 3E).

**Figure 3:**
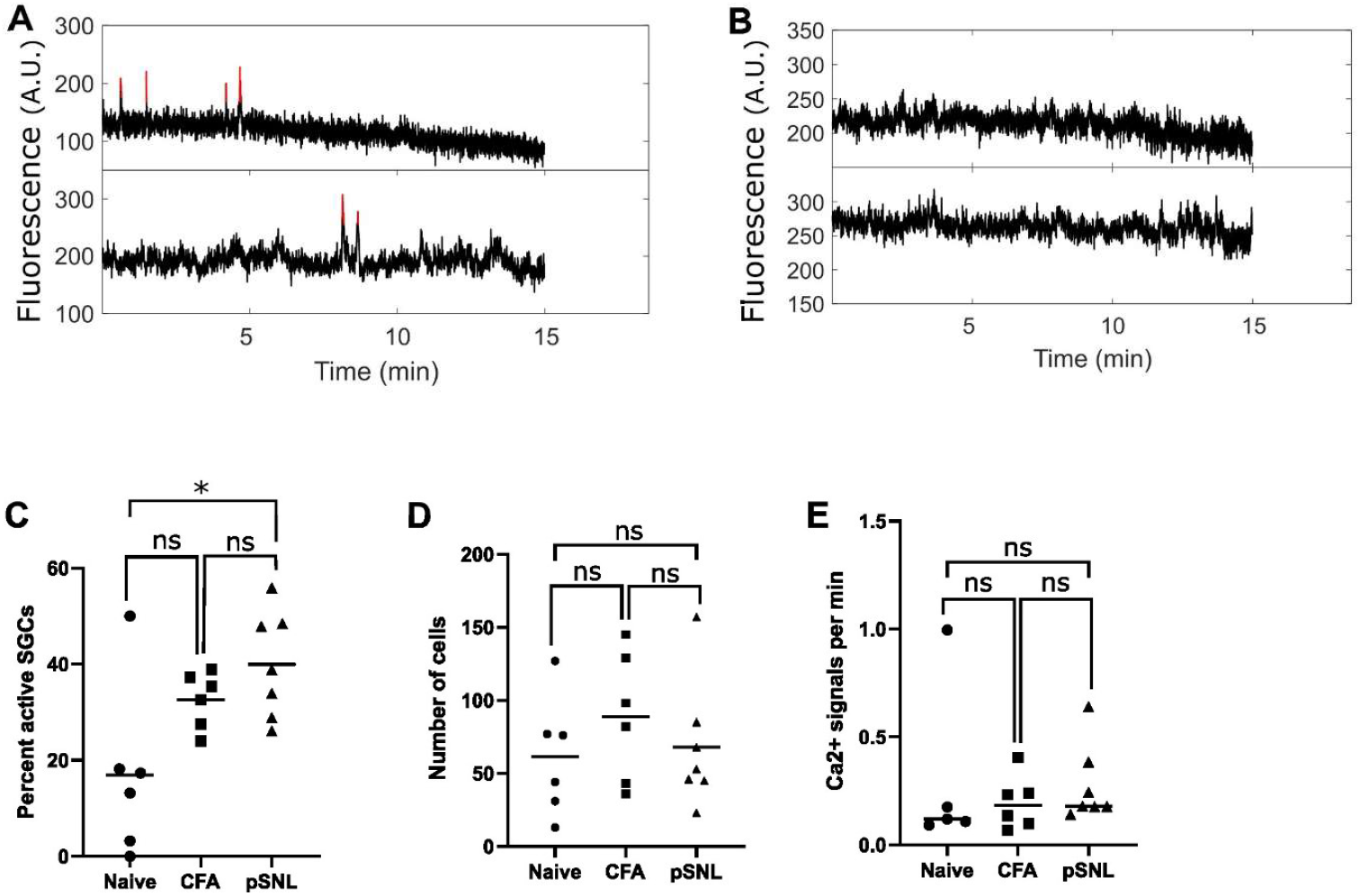
Quantification of Ca^2+^ responses in SGCs after peripheral sensitization. **(A)** Two example traces of active SGCs in the baseline condition, i.e. without stimulation. The red lines indicate responses above background noise **(B)** Two example traces of inactive SGCs. **(C)** Percent SGCs that have at least one Ca^2+^ response in each mouse across the conditions: naïve (n=6 mice), CFA (n=6 mice) & pSNL (n=7 mice). Ordinary one-way ANOVA test followed by Tukey’s multiple comparisons test of means comparing Naïve vs CFA (p=0.114), Naïve vs pSNL (*p=0.01) and CFA vs pSNL (p=0.55). **(D)** Number of SGCs imaged in each mouse across the three conditions. Ordinary one-way ANOVA test followed by Tukey’s multiple comparisons test of means comparing Naïve vs CFA (p=0.52), Naïve vs pSNL (p=0.96) and CFA vs pSNL (p=0.67). **(E)** Averaged number of Ca^2+^ signals per minute across active SGCs within each mouse. SGCs with at least one Ca^2+^ response were considered active and thus included in this plot. Ordinary one-way ANOVA test followed by Tukey’s multiple comparison test of means comparing Naïve vs CFA (p=0.77), Naïve vs pSNL (p=0.99) and CFA vs pSNL (p=0.81).

After *in vivo* imaging, DRGs from the pSNL animals were used to quantify how many of the SGCs were in fact surrounding injured neurons. To perform this analysis DRGs were immersion-fixed in 4% PFA and stained with an antibody against the injury marker ATF3 (Fig. 4A&B). We observed that slightly less than half of the SGCs that we recorded from in pSNL animals surrounded injured neurons (mean = 38%), while the remaining 62% surrounded uninjured neurons (Fig. 4C). This means that the size of the effects plotted in Fig. 3C may be an underestimate, since our data include many SGCs that were not in the proximity of sensitised neurons.

**Figure 4:**
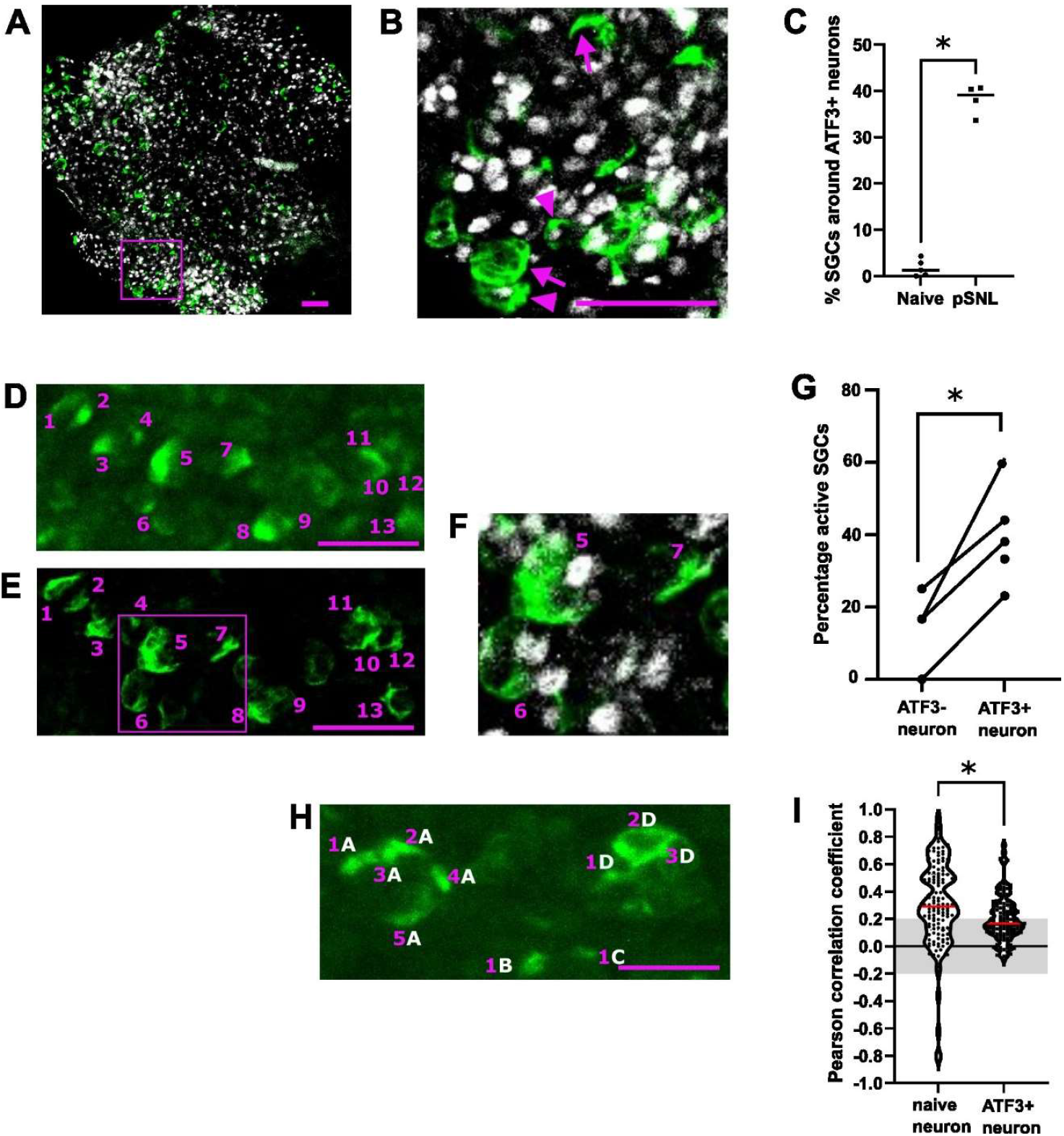
Analysis of SGCs surrounding injured or uninjured neurons after nerve ligation. **(A**) Representative image of immunostained L4 DRG, following in vivo Ca^2+^imaging. The tissue was post-fixed by immersion and stained with the injury marker ATF3 (grey) and GFP to enhance GCaMP6 signal (green). Scale bar = 100μm. **(B)** Magnified view of the region highlighted within the purple box in (A). SGCs surrounding uninjured (ATF3-) vs. injured (ATF3+) neurons are highlighted by arrows vs. arrowheads, respectively. Scale bar = 100μm. **(C)** Quantification of percentages of SGCs surrounding ATF3+ neurons in naïve (n = 5) and pSNL conditions (n = 4). Each datapoint show the quantification from one immunostained L4 DRG after in vivo Ca^2+^ imaging. Mann-Whitney test, *p = 0.016. **(D)** Representative image of SGCs during in vivo Ca^2+^ imaging recording with SGCs shown in green. The recorded SGCs are labelled with purple numbers. Scale bar = 100 μm. **(E)** Confocal image of whole mount DRG stained against GFP to enhance GCaMP6 signal (green) in SGCs following imaging. SGCs were matched to their in vivo Ca^2+^ imaging profile, i.e. purple cells 1-13 imaged via confocal in (E) are the same as purple cells 1-13 in (D), recorded during an imaging session. Scale bar = 100 μm. **(F)** Magnified view of the region highlighted with purple in (E). The magnification highlights SGC numbers 5-7 including the immunostaining of the injury marker ATF3 (grey) to identify which SGCs surround injured neurons. SGC 5 surrounds an ATF3+ neuron, SGC 6 is undetermined and SGC 7 surrounds an ATF3-neuron. **(G)** The percentage of active SGCs surrounding either ATF3+ or ATF3-neurons after nerve injury. Data derived from n = 4 videos from 3 mice; paired t-test; *p = 0.019. Each set of paired datapoints show the quantified percentage of active SGCs around either ATF3+ or ATF3-neurons in each video. In the fourth pSNL mouse plotted in C, no SGCs surrounding ATF3-neurons were identified in the corresponding video. It could therefore not be included in the analysis. **(H)** Representative image of SGCs during in vivo Ca^2+^ recording with SGCs shown in green. The SGC-neuron units are marked with white letters, and individual SGCs in each unit are marked with purple numbers. The individual SGCs are identified with Suite2p based on their Ca^2+^ activity. Scale bar = 100 μm. **(I)** The recorded Ca^2+^ fluorescent traces from SGCs in the same unit were compared with Pearson’s correlation. In the graph, SGCs from the pSNL group are only included if they surround an ATF3+ neuron. For the SGCs surrounding ATF3-neurons, it was only possible to perform six comparisons. Instead, SGC from naïve mice were analysed. The y-axis shows the correlation coefficient R. Traces with an R between -0.2 and 0.2 were considered to be uncorrelated (grey shaded area). Data derived from n = 124 comparisons from the naïve group and n = 133 from the ATF3+ group. Each datapoint represents the correlation between two Ca^2+^ traces from SGCs surrounding the same neuron. Statistical analysis was performed using a non-parametric, two-tailed Mann Whitney test, *p value > 0.0001.

To address this, we used careful alignment to overlap the *in vivo* calcium signals we observed with the ATF3 labelling post-mortem. By combining these two imaging modalities, we were able to compare the activity of SGCs surrounding uninjured vs. injured neurons in the same DRG (Fig. 4D-F). We found that the % of active SGCs surrounding injured neurons was larger than that surrounding uninjured neurons (d = 2, n = 4 videos, Fig. 4G). As expected from this within-animal comparison, the effect was slightly larger than that observed in our less sensitive, between-animal comparison (Fig. 3C).

To investigate whether we could detect a rise in intracellular calcium waves exchanged between SGCs surrounding the same injured neuron, we assessed whether they are more correlated in their activity compared to SGCs which surround the same naïve neuron. First, we identified SGCs surrounding the same neuron (Fig. 4H) and whether they were surrounding an injured or uninjured neuron based on neuronal ATF3 expression. Next, we calculated the Pearson correlation of the calcium traces from SGCs surrounding the same neuron. Our data did not provide any evidence for a rise in intracellular calcium waves in the injured condition, thus whether we examined groups of SGCs around injured or naïve neurons – their activity was not any more or less correlated with each other. The spread of correlations ranged from 0.2 to 0.9 in both conditions, with the mean correlation actually higher for SGCs surrounding naive rather than injured neurons (Fig. 4I).

## Discussion

Our study set out to use a transgenic mouse model to image SGC calcium transients *in vivo*. Our results are the first to show in a whole-animal physiological setting that there is more SGC activity in a sensitized peripheral nervous system. Specifically, an increased percentage of active SGCs could be observed in inflammation (CFA) and nerve injury (pSNL) models compared to naïve controls.

Moreover, a higher percentage of SGC-generated calcium transients could be observed around ATF3+ injured neurons, compared to their uninjured, ATF3-counterparts. However, with the exception of repeated suprathreshold stimuli, the response of individual SGCs did not appear to be time-locked with actual neuronal firing.

The general lack of time-locked SGC calcium transients suggests that they are not directly induced by neuronal action potentials. This finding is surprising, since it conflicts with prior *in vitro* work ^21–23^. For example, it has been shown by Zhang et al. that cultured DRG neurons release ATP from their soma upon firing, while the SGCs in turn detect the ATP and respond with an intracellular calcium signal ^21^. One reason for this discrepancy could be that the neurons in the culture are electrically activated for 30 seconds, while we did electrical pulses for no longer than 10 seconds. This explanation is also supported by the observation that we do see some time-locked responses in the injured condition with repeated suprathreshold stimuli, which is an extraordinary physiological situation.

The *in vivo* system we used in this study did have some challenges. Specifically, the transgenic mouse model has sparse labelling of SGCs, and the neuronal cell bodies in the imaged DRG do not all project to the sciatic nerve ^42^. In theory, these two challenges could mean that we have missed time-locked signals. However, we think this is unlikely, since we were able to activate approximately 75% of labelled neurons with our electrical stimulation paradigm (437 neurons out of 575 labelled neurons from 2 mice, Suppl. Fig 1), and have imaged over 1378 SGCs from a total of 19 mice. Currently, we therefore cautiously conclude that neuronal-SGC signalling with Ca^2+^ as second messenger may be less important in homeostatic conditions than we would have previously thought.

During peripheral sensitization, we observe an overall, diffuse increase in SGC Ca^2+^ activity. The function of this is currently unknown. In general, intracellular Ca^2+^ concentrations are used as second messengers and can influence a broad variety of signalling pathways. We speculate that in this case, the increase in Ca^2+^ activity could be a reflection on the need for more buffering of the extracellular space in the SGC-neuron unit. This might be necessary to accommodate any extra ions and neurotransmitters released into the SGC-neuron unit, as a result of the increased neuronal activity known to occur after inflammation or nerve injury ^43^. In line with this, it has recently been suggested by Schulte et al. ^44^ that SGCs undergo changes in metabolic function after nerve injury, evidenced by a change in marker expression of GFAP and GS.

Like our team, Chen et al. ^45^ were also able to analyse calcium signals in SGCs. They did not analyse them at single cell level, but examined aggregated calcium signal across a wider area, reporting an increase when ATP was applied directly to the DRG. While Chen et al. did not investigate pain models, their results raise the possibility that the increase in activity we found after inflammation or nerve injury might be linked to ATP released by injured neurons.

It should be noted that our model has some limitations. Firstly, we focused our attention on calcium responses in SGCs, but there are many other ways in which these cells might functionally respond to neuronal activity, such as release of messenger molecules. Secondly, our setup does not permit a distinction between increased SGC-SGC communication (e.g. via gap junctions) vs. increased communication between one neuron and each of its SGCs individually. Thirdly, labelling of SGCs using the *Fabp7*-CreER mouse line proved extremely sparse and variable. This meant that the results described above were very time-consuming to obtain, with every other animal labelled too poorly to proceed to imaging. Moreover, even in the animals that we could image, many SGCs would have remained unlabelled, which means that we would have missed other smaller effects in addition to the large changes we observed. Furthermore, our imaging was done with mice under anaesthesia which might dampen their responses as e.g. reported in astrocytes ^46^.

In conclusion, *in vivo* calcium imaging of SGCs has permitted us to visualise some of the function of these cells in intact but anaesthetised mice, during health and disease. It appears that they work according to their own schedule, but crucially, increase their overall output in more metabolically demanding settings like inflammation and nerve injury. At present, it is not clear whether this increase in activity is maladaptive or not. Currently not much is known about the downstream effects of Ca2+ signalling in SGCs, but their altered function in disease states is generally believed to enhance neuronal excitability and pain ^47^. The additional *in vivo* data we provide here thus strengthen the case for targeting SGC function pharmacologically, in order to bolster neuronal function and alleviate pain in neuropathy.

## Materials and Methods

### Transgenic animals for investigating SGCs

All mice were housed under standard conditions with 12hr light/dark cycles and free access to chow and water. To create mice with transgene expression in SGCs the Cre-dependent mouse strains flox-STOP-*GCaMP6s* (The Jackson laboratory, Bar Harbor, ME, 028866) and the nuclear-tagged reporter line flox-STOP-*Sun1GFP* (The Jackson laboratory, #030958) were crossed with the *Fabp7*-creER mouse strain (BRC Riken, Ibaraki, Japan, #RBRC10697) ^33^. The mice were bred as homozygotes for either GCaMP6 or Sun1GFP and hemizygote for *Fabp7*-CreER. The genotypes of the mice were confirmed using standard PCR conditions with the primers listed in Table 2. For expression of GCaMP6s or GFP the mice were given 180mg/ml tamoxifen (Merck KGaA, Darmstadt, Germany, #T5648-1G) dissolved in corn oil (Merck, #C8267-500ML) with oral gavage for five consecutive days. We found this to be the most effective tamoxifen administration protocol, having also tried delivering tamoxifen via food and via serial intraperitoneal injections. No protocol we tried enabled us to achieve consistent transgene expression. Two weeks were allowed between the tamoxifen treatment and further procedures such as nerve ligation and *in vivo* Ca^2+^ imaging. All mice used in the study was between 8-17 weeks of age. All animal experiments were performed in accordance with the United Kingdom Home Office Legislation (Scientific Procedures Act, 1986) and were approved by the Home Office to be conducted at King’s College London under project license number P57A189DF.

**Table 2:**
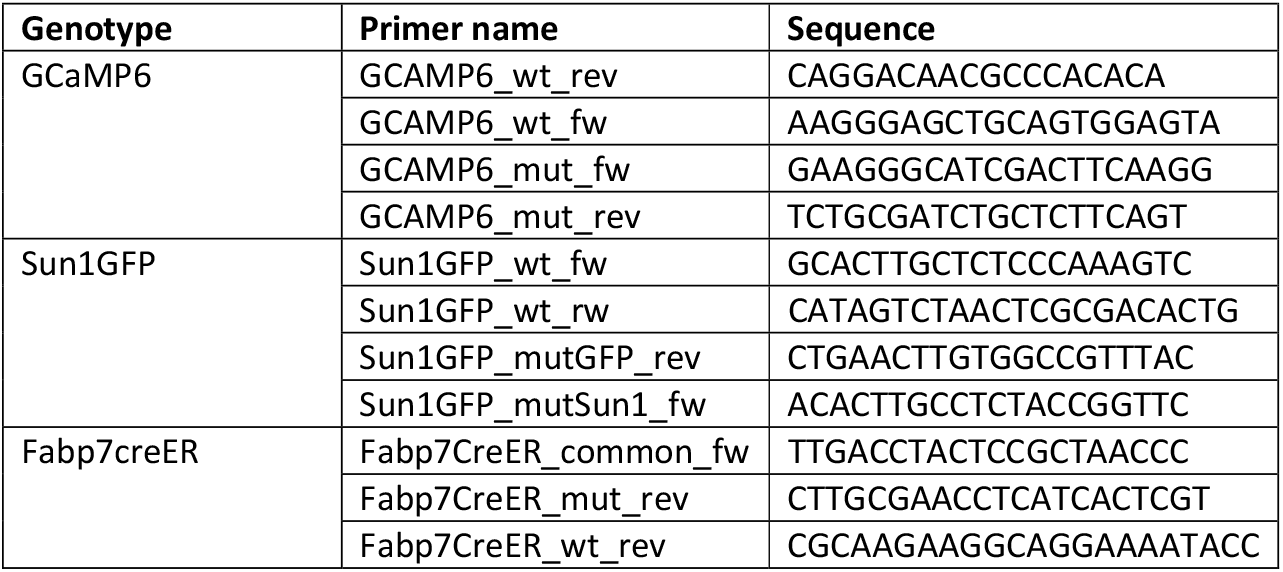
Primers used for genotyping. Annealing temperature at 59°C for all reactions.

### Immunohistochemistry of DRGs

DRGs from tamoxifen treated *Fabp7*-CreER-*GCaMP6s* mice were dissected following transcardial perfusion with PBS and 4% PFA. The DRGs were post-fixed for 2 hrs in 4% PFA before they were washed in PBS and incubated overnight in 30% sucrose for cryoprotection. The tissue was embedded in OCT (CellPath Ltd, Newton, UK, #KMA-0100-00A) and stored at -80°C. The embedded tissue was cut at 10 μm on a cryostat and collected on Superfrost plus slides (ThermoFisher Scientific, #J1800AMNZ). The slides were baked for 30 min at 50°C to ensure adhesion of the tissue sections before the OCT was washed off with PBS. A hydrophobic barrier was drawn (Vector laboratories, Newark, CA, #H-4000) around the tissue sections. When the barrier was dry, the slides were washed in PBS with 0.3% triton x-100 (Santa Cruz Biotechnology Inc, Dallas, TX, #sc-29112A), and the tissue sections were incubated with blocking solution (10% donkey serum, 0.3% triton x-100 in PBS) for at least 1 hr in a humified chamber. Next, they were incubated in primary antibody mix (see Table 3) diluted in blocking solution overnight at room temperature. The following day, the tissue sections were washed 3x5 min in PBS with 0.3% triton x-100 before they were incubated in secondary antibody mix (see Table 4) diluted in blocking buffer for 4 hrs at room temperature protected from light. The slides were washed 3x5 min in PBS with 0.3% triton x-100 before they were coverslipped using mounting medium containing DAPI (Invitrogen, #00-4959-52). The slides were imaged with a Zeiss LSM 710 confocal microscope and quantified using ImageJ. For any given experiment, 4-6 sections were stained from each mouse. The stained cells were quantified and reported as absolute counts or percentages of total amount of cells as determined based on number of nuclei. For each staining, tissue from 6-7 mice was used, enough to detect any sizable recombination events outside of our target cell type. For example, a one-sample t-test with n = 6 would give an 80% chance to detect significant recombination in a cell type other than SGCs (d = 1.79, i.e. a change from 0% to 4% recombination with a standard deviation of the mean difference as large as 5).

**Table 3:** Primary antibodies used for immunohistochemistry (IHC), DRG whole mount (whole mount) or flow cytometry (flow). The last antibody in the table was conjugated in house with APC. The catalogue number for the conjugation kit is added in the table below the catalogue number for the antibody.

| Application | Antibody | Dilution | Conjugation | Catalogue # | Supplier |
| --- | --- | --- | --- | --- | --- |
| IHC | Rabbit α-GS | 1:5000 | no | Ab49873 | Abcam |
| IHC | Chicken α-MPZ | 1:250 | no | Ab39375 | Abcam |
| IHC | Rabbit α-GFP | 1:250 | no | A11122 | Invitrogen |
| IHC | Goat α-Fabp7 | 1:500 | no | AF3166 | R&D |
| IHC/whole mount | Chicken α-GFP | 1:5000 | no | Ab13970 | Abcam |
| Whole mount | Rabbit α-ATF3 | 1:500 | no | NBP1-85816 | Novus |
| Flow | α-CD31 | 1:300 | BV711 | 740680 | BD Biosciences |
| Flow | α-CD11b | 1:50 | PE | 101208 | Biolegend |
| Flow | α-CD45 | 1:300 | PerCP Cy5.5 | 45-0451-80 | eBiosciences |
| Flow | Rabbit α-GS | 1:200 | APC | Ab49873<br>Ab201807 | Abcam<br>Abcam |

**Table 4:**
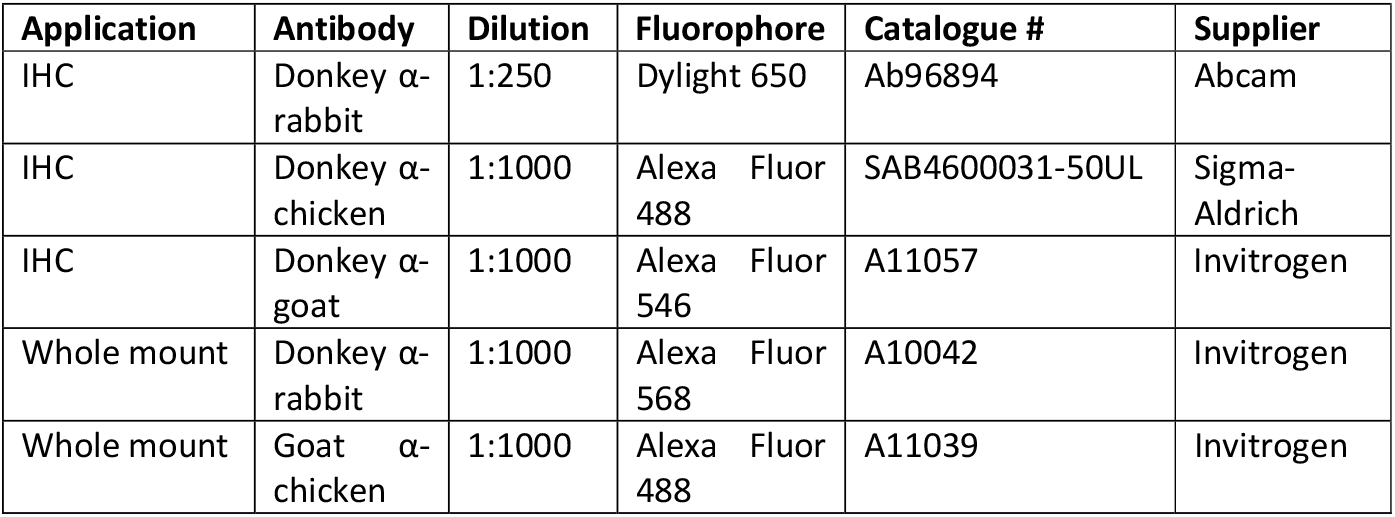
Secondary antibodies used for immunohistochemistry (IHC) or DRG whole mount (whole mount).

### Flow cytometry of DRGs

L3-L5 DRGs from tamoxifen treated *Fabp7*-CreER-*Sun1GFP* mice were dissected following transcardial perfusion with PBS ^34^. The DRGs were enzymatically dissociated by incubation with digestion mix containing F12 with 3mg/mL dispase II (Sigma Aldrich, #04942078001), 12.5 mg/mL collagenase type IA (Sigma Aldrich, #C9891) and 10 mg/mL DNase I (Sigma Aldrich # 10104159001) for 45 min at 37°C. The cells were triturated with a p1000 pipette tip until homogenous and filtered through a 70μm cell strainer into an Eppendorf tube. They were spun down at 200xg for 5min and resuspended in Hank’s balanced salt solution (HBSS) (Gibco, New York, NY, #14175095). Next, the cells were moved to a 96 well plate and protected from light thereon. Staining proceeded with 50μL Near-IR live/dead (Invitrogen, Waltham, MA #L34975) diluted 1:1500 in HBSS. After a 30min incubation on ice, we added 50 μL of primary antibody mix (see Table 3) diluted in FACS buffer containing HBSS with 0.4% bovine serum albumin (Sigma-Aldrich, #A3983), 15mM HEPES (Gibco, #15630080), and 2 mM EDTA (Invitrogen, #15575-038). This was left for a further 30min on ice, before cells were spun down at 200xg for 5 min and fixed in 4% PFA for 10 min followed by permeabilization in perm buffer (Invitrogen, #00-8333-56) with 10 μg/mL Hoechst (ThermoFisher Scientific, Waltham, MA, #62249) for 5 min at room temperature. To perform the intracellular staining the cells were incubated with the GS antibody (see Table 3) diluted in perm buffer for 30 min on ice. Finally, the cells were spun down at 200xg for 5 min and resuspended in FACS buffer followed by flow cytometric analysis at the NIHR BRC flow core facility at King’s College London on a BD Fortessa or BD FACSCanto. Fluorescence minus one (FMO) stainings were used as staining controls. For each flow cytometric analysis, tissue from 5 mice was used, which powered us to detect significant off-target recombination (80% chance to detect d = 2.02 with a one samples t-test and alpha = 0.05, i.e. a change of 0% to 4.25% recombination).

### Neuropathic pain model

Partial sciatic nerve ligation (pSNL) was performed in the left hindleg using a paradigm originally developed in the rat ^35^. In short, mice were administered 0.05 mg/kg of the analgesic buprenorphine (Ceva, Libourne, France, #Vetergesic) and anaesthetised with 2% isoflurane (Henry Schein, Melville, NY, #988-3245). After shaving the fur covering the left hip and leg region, a skin incision was made, and the sciatic nerve was exposed by blunt dissection of the muscle layer. Approximately 2/3 of the sciatic nerve was tightly ligated with a 5.0 suture (Ethicon, Cincinnati, OH, W9982). The incision was closed with a surgical clip (World Precision instruments, Sarasota, FL, #500433), before mice were placed in a clean cage and left to recover in the animal unit. The weight of the mice was monitored daily until the pre-procedure weight was re-established. Mice were used for imaging experiments at day seven after the procedure.

### Inflammatory pain model

Mice were anaesthetised with 2% isoflurane and injected with 20 ul complete Freund’s adjuvant (CFA), 1mg/ml (Sigma, #F5881-10ML) in the left hind paw. They were used the following day for *in vivo* Ca^2+^ imaging experiments.

### *In vivo* Ca^2+^ imaging

Terminal *in vivo* Ca^2+^ imaging was performed on SGCs in *Fabp7*-CreER-*GCaMP6s* mice and on neurons in wild-type mice labelled through viral transduction of GCaMP6s. For the latter, 5μl of AAV9 pAAV-Syn-GCaMP6s-WPRE.SV40 virus (Addgene, Watertown, MA, #100843-AAV9) was injected into C57BL/6J pups 3-6 days after birth as previously described ^36^.

For imaging, anaesthesia was induced with an i.p. injection of 12.5% w/v urethane (Merck, U2500-100G) and surgical depth was achieved using ∼2% isoflurane. The exact level of isoflurane was determined based on hindlimb and corneal reflex activity. The core body temperature was maintained close to 37°C using a homeothermic heating map with a rectal probe (FHC, Bowdoin, ME). An incision was made in the skin on the back, and the muscle overlying L3, L4 and L5 DRG was removed. The L4 DRG was identified based on the position of the hipbones and the vertebra. The bone covering the L4 DRG was carefully removed, and the underlying epineurium and dura mater over the DRG were washed and kept moist with saline. The exposed L4 DRG was covered with silicone elastomer (World Precision Instruments, #Kwik-sil) to avoid drying and maintain a physiological environment as described previously ^20^. The mouse was then placed under an Eclipse Ni-E FN upright confocal/multiphoton microscope (Nikon, Tokyo, Japan). All images were acquired using a 20x dry objective and a 488-nm Argon ion laser line. Time series recordings were made with a resolution of 512x512 pixels and a fully open pinhole. Image acquisition was at either 1.8 or 3.6 Hz depending on experimental requirements. When possible, up to three distinct areas of the DRG were imaged for each mouse to record from as many SGCs as possible. At the end of the experiment the mouse was culled with an intracardial injection of pentobarbital (Euthatal; Merial, Duluth, GA, #P02601A). Data were collected from 6 naïve mice (two male and four female), 6 CFA treated mice (one male and five female) and 7 pSNL mice (two male and five female) between 8 to 17 weeks of age. We chose these sample sizes as we were looking for large effects between biological replicates, and with *n* = 6 one has an 80% chance to detect f = 0.89 with alpha = 0.05 using a one-way ANOVA, as we did. Of note, these considerations are at their most conservative, as we are not taking into account the large number of cells we are recording from in each mouse. Due to the nature of these experiments it was not possible for the experimenter to be blind to treatment condition when collecting the data.

### Activation of neurons during *in vivo* Ca^2+^ imaging

To activate neurons in the L4 DRG the sciatic nerve was electrically stimulated using a custom-made cuff electrode with insulated steel wire as previously described ^20^. Pulses of 500 μA amplitude and a duration of 400 μsec were used to activate A fibres in the sciatic nerve, while suprathreshold stimulation with pulses of 1 mA amplitude and a duration of 1 ms were used to activate both A fibres and C fibres ^20^ (Suppl. Fig 1). Furthermore, neurons in the sciatic nerve were activated by squeezing the hind paw with a pair of tweezers for a duration of 5 sec.

### Analysis of *in vivo* Ca^2+^ imaging data

Drift in the time series recordings was corrected using the registration framework in Suite2p v0.10.3^37^. To extract the Ca^2+^ traces, regions of interests (ROIs) corresponding to individual SGCs were identified using Suite2p together with Cellpose v1.0 ^38^ (Suppl. Fig 2). The detected ROIs were quality checked manually leading to exclusion of ROIs on the edge of the video and the addition of extra ROIs when needed.

The output from Suite2p (‘Fall.mat’) was modified with a custom matlab script which dynamically subtracts the background neuropil fluorescence, i.e. signal in the pixels surrounding each ROI. Next, the time series recordings were analysed to identify the Ca^2+^ responses. The threshold for a positive response was taken as a signal above 25% of the mean fluorescent signal for the full video (or until pentobarbital administration) plus 2 * standard deviation. For 13 ROIs (four in naïve condition, four in CFA condition and five in pSNL condition) the mean was too high to be used as baseline because of a very high level of Ca^2+^ activity throughout the recording. In these cases, a baseline interval without activity was identified individually for each ROI. Responses only spanning one frame were determined to be noise and were thus removed from further analysis. Since our computational pipelines are largely automated, these analyses were not generated by an experimenter blind to treatment condition.

Visualization with raster plots were generated in Matlab using plotSpikeRaster_v1.2 ^39^, after the data were changed to a binary format based on whether the fluorescent signal reached the threshold or not.

The frequency of activity was calculated for each active SGCs in a mouse as the number of Ca^2+^ responses per minute. A SGC was considered active, and thus included in this analysis, if it had at least one Ca^2+^ response during the imaging session.

A Ca^2+^ response was determined to be time-locked if it was initiated within 7.5 seconds of the noted time for the stimulation of the neurons. Thus, the chance for one cell to appear time-locked was calculated as: FB = R * 7.5 sec, where FB is the number of background firing events in every 7.5 second interval and R is the average firing rate of cells per second. To determine the number of

time-locked responses expected by chance, we multiplied the background firing events of one cell (per 7.5 second interval) with the number of cells in a given dataset.

Correlation analysis was performed to investigate if SGCs surrounding individual injured neurons are more correlated than SGCs surrounding individual uninjured neurons. The ROIs identifying SGCs surrounding the same neurons were identified manually, and their fluorescent traces were compared with Pearson’s correlation in matlab.

### Whole mount staining of DRGs

Following *in vivo* Ca^2+^ imaging, DRG were dissected and immersion fixed in 4% PFA for at least 3 hrs. They were then washed in PBS, followed by incubation in blocking solution (10% donkey serum and 0.3% triton x-100 in PBS) for at least 1 hr. Next the DRGs were stained with primary antibody mix (see Table 3) diluted in blocking solution for 3 days at room temperature and protected from light. They were then washed 3x5min in PBS with 0.3% Triton x-100, before being incubated with secondary antibody mix (see Table 4) diluted in blocking solution for 2 days at room temperature and protected from light. The DRGs were imaged in two rounds to increase the chance of identifying the same SGCs visualised during *in vivo* Ca^2+^ imaging. First, we used a fluorescent microscope (Zeiss Axio Imager Z1) covered only in PBS to retain as much original 3D structure as possible. Next, the DRGs were covered with mounting medium and a glass slide prior to imaging on a Zeiss LSM 710 confocal microscope with Z-stacks. The images were used to identify which SGCs in the *in vivo* Ca^2+^ imaging data were surrounding injured neurons in the pSNL condition. Great care was taken to avoid false negatives, i.e. a neuron was only identified as uninjured if it had been imaged in its entirety in the z-plane, ensuring that no ATF3+ signal in the nucleus could have been missed. The analysis was performed by an investigator blind to treatment condition and fluorescent traces.

### Statistical analysis

All statistical analyses were performed in GraphPad Prism 9. For comparisons between the three groups (Naïve, CFA and pSNL) ordinary one-way ANOVA followed by Tukey’s multiple comparisons test were used. For the comparison between the percentage of SGCs surrounding ATF3 positive neurons in naïve versus pSNL animals a two-tailed Mann-Whitney test was used. For the comparison between percent active SGCs surrounding injured or uninjured neurons a paired t-test was used.

Finally for the comparison of the distribution of the Pearson correlations a non-parametric two-tailed Mann-Whitney test was used.

## Supporting information

Video1_naive

Video2_CFA

Video3_pSNL

## Data availability

Example videos of *in vivo* calcium imaging, raw fluorescence traces from Suite2p and matlab scripts are made available here: https://osf.io/jg5zr/?view_only

## Acknowledgements

This research was supported by a Lundbeck Foundation Fellowship (R293-2018-960) and a grant from IMK Almene Fond (30206-372) for S.E.J.. F.D. has received funding through an MRC New Investigator Research Grant (MR/P010814/1) and a Medical Research Foundation Prize (MRF-160-0015-ELP-DENK-C0844). G.G. is funded by an Advanced Pain Discovery Platform UKRI MRC grant (MR/W027518/1). K.I.C. is funded by an Anne McLaren Fellowship, University of Nottingham.

The research has also received support from the National Institute for Health and Care Research (NIHR) Biomedical Research Centre based at Guy’s and St Thomas’ NHS Foundation Trust and King’s College London. The views expressed are those of the authors and not necessarily those of the NHS, the NIHR or the Department of Health and Social Care. This research was funded in whole or in part by the UK Research & Innovation.

For the purpose of Open Access, the author has applied a CC BY public copyright licence to any Author Accepted Manuscript (AAM) version arising from this submission.

## Supplementary figures

**Supplementary Figure 1:**
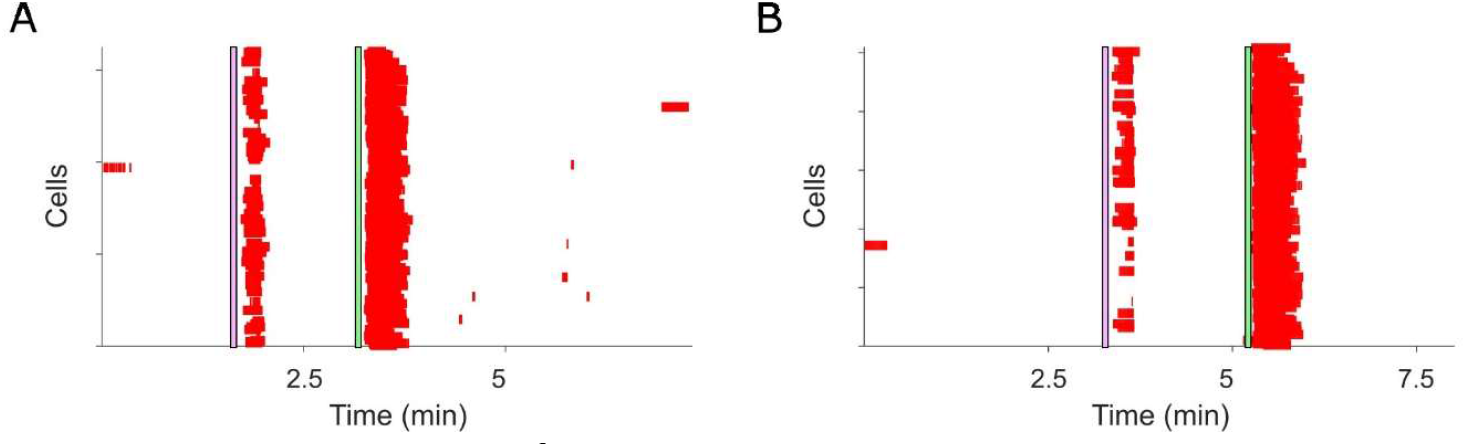
In vivo Ca^2+^ imaging of neurons. **(A)-(B)** Raster plots showing neuronal activation following electrical stimulation. Each raster plot represents data from one mouse. The red annotations indicate significant Ca^2+^ signal above baseline. Coloured lines (pink and green) indicate times at which neurons were stimulated: A fibre stimulation for 10sec at 4 Hz (light pink), suprathreshold stimulation for 10sec at 4 Hz (light green).

**Supplementary Figure 2:**
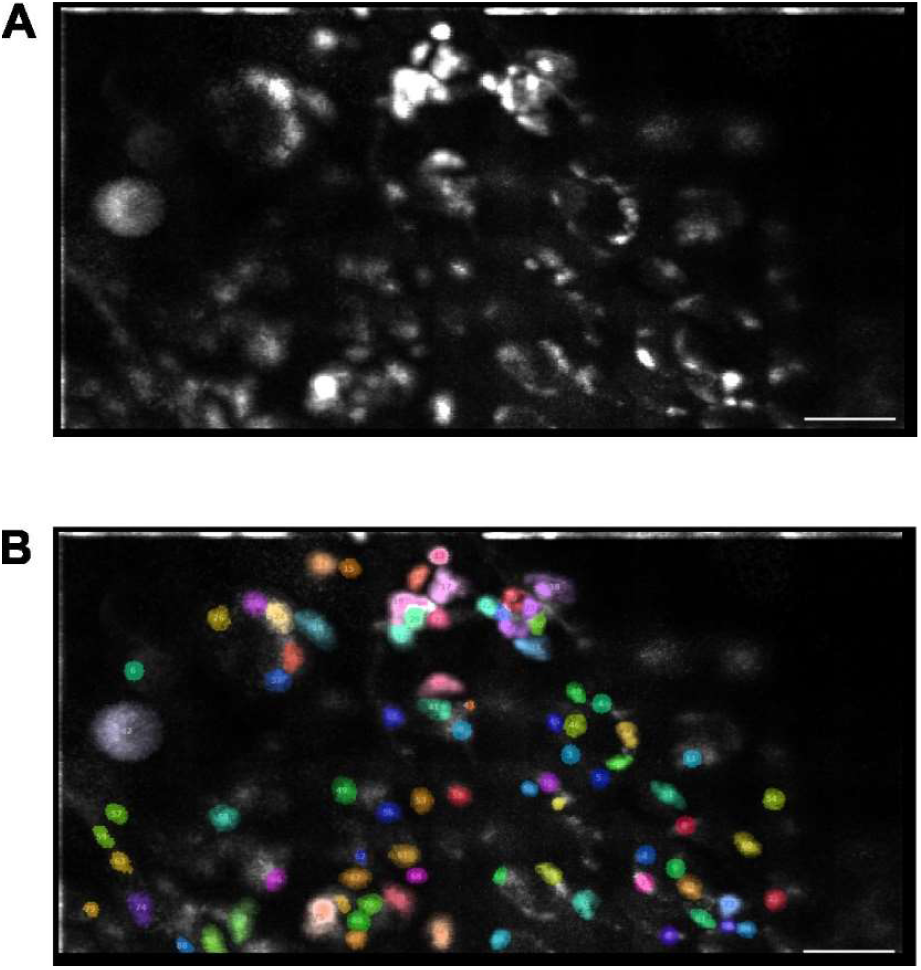
Identification of individual SGCs with Suite2p and Cellpose. **(A)** Example of max projection of GCaMP6 signal in SGCs from in vivo Ca^2+^ imaging experiment. Scale bar = 100μm **(B)** Detected regions of interest (ROI) and ROI number overlaid on the max projection. Suite2p and Cellpose take the activity throughout the video into account when annotating the ROIs, which ensures that individual SGCs can be distinguished when they fill with calcium at different times, either throughout the recording or as they die at the end. Scale bar = 100μm

**Supplementary Figure 3:**
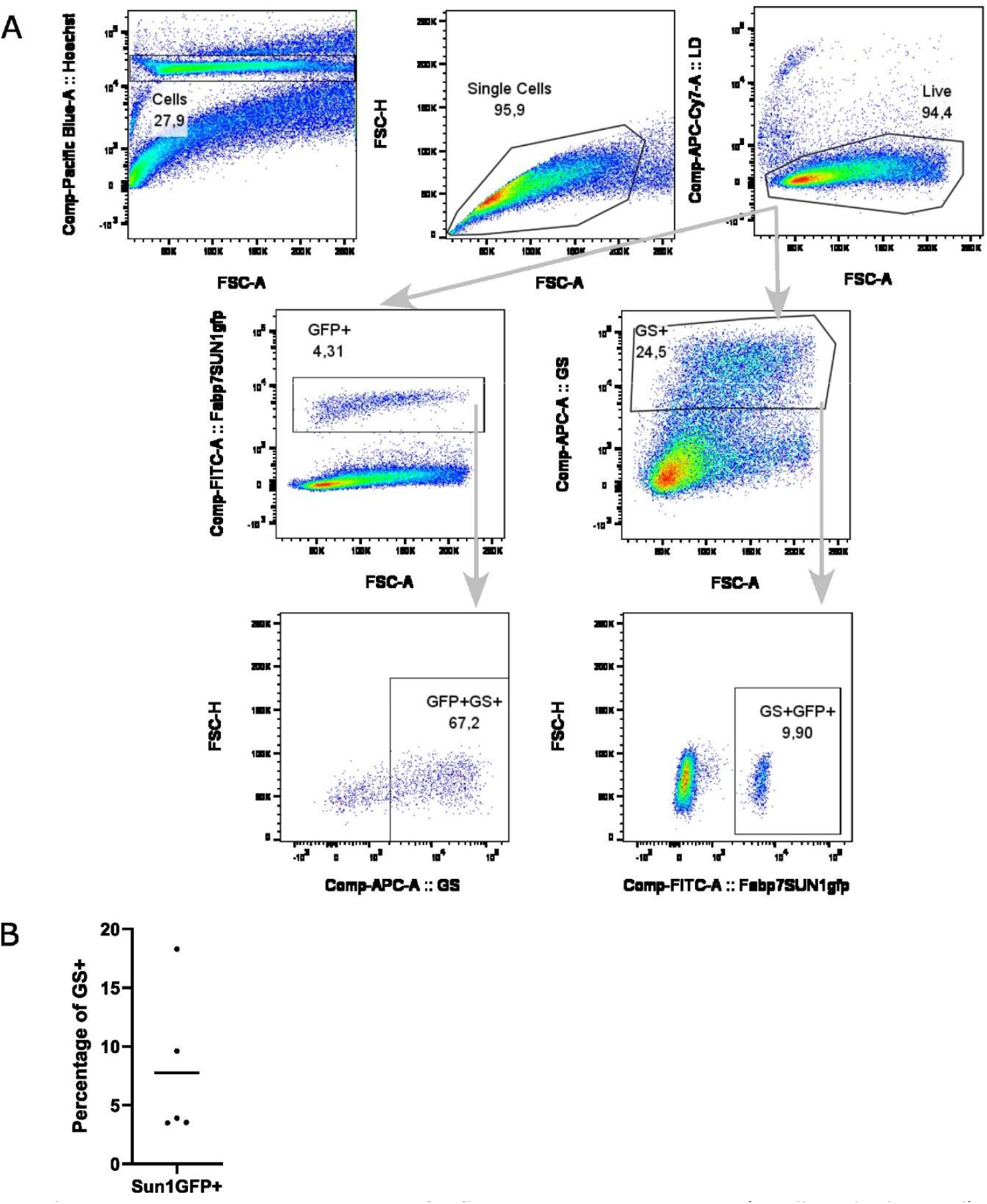
Gating strategy for flow cytometric experiment (small antibody panel) **(A)** Gating strategy for Figure 1D. Samples run on a BD FACSCanto. First, cells are distinguished from debris with Hoechst staining. Next, single cells followed by live cells are identified. From here either the Fabp7-CreER-Sun1GFP positive cells or the GS positive cells are identified and lastly the percentage of Sun1GFP and GS double positive cells are determined. All gates are placed based on fluorescence-minus-one (FMO) controls. **(B)** Quantification of the percentage of single, live and GS positive cells that are also positive for Sun1GFP. The black line represents the mean percentage, n=5 mice.

**Supplementary Figure 4:**
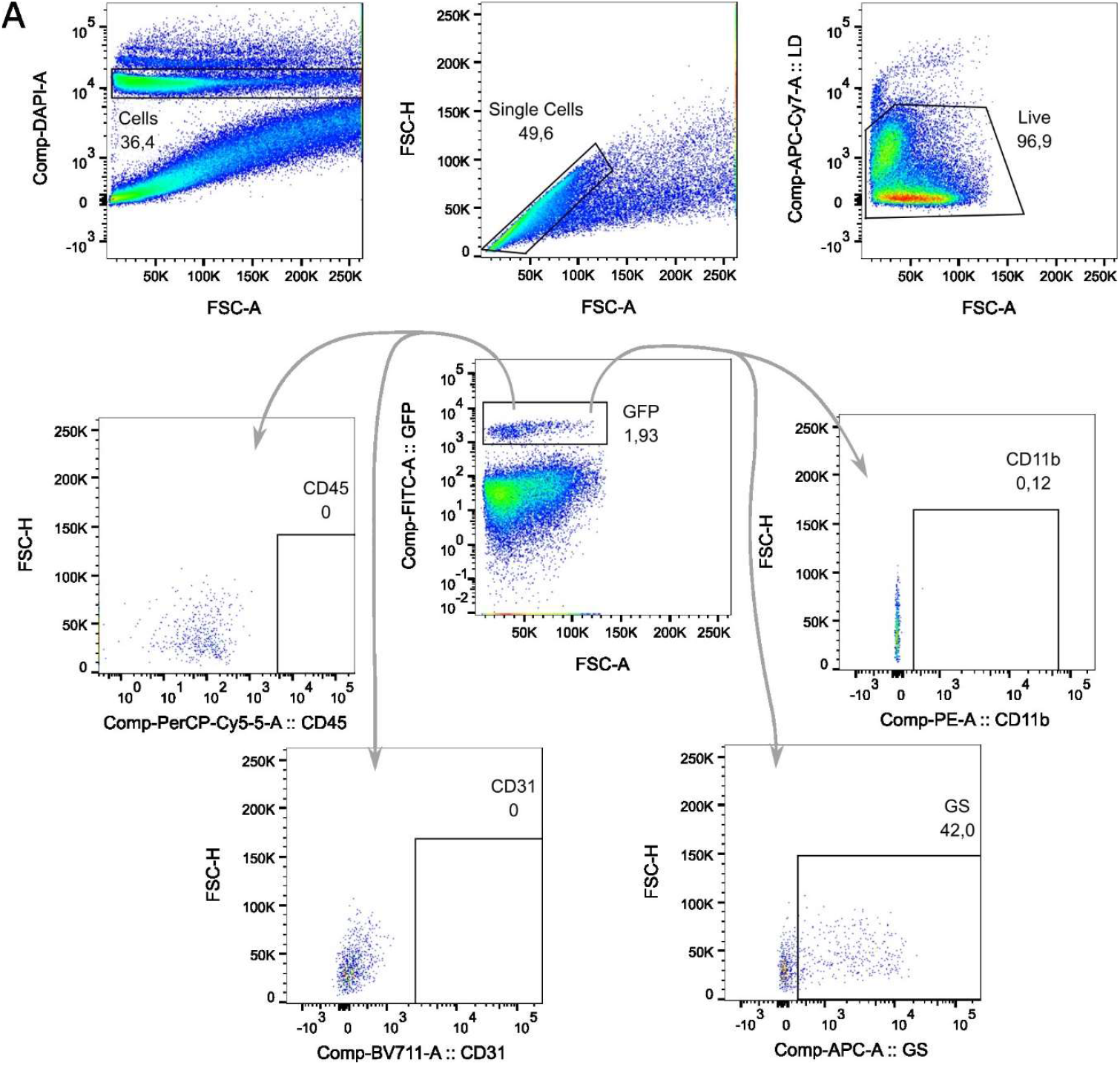
Gating strategy for flow cytometric experiment (big antibody panel) **(A)** Gating strategy for Figure 1H. Samples run on a BD Fortessa. In the first three plots Hoechst positive, singlets and live cells are identified. Next, Fabp7-CreER-Sun1GFP positive cells are identified. From there, the percentage of GFP+ cells also positive for the cell markers CD11b, CD45, CD31 and GS are determined. All gates are placed based on FMO controls.

## Supplementary videos

Supplementary videos have been uploaded as supplementary material.

Furthermore, the videos can be downloaded from the following links. Open the link and use the small menu (button with three small dots) on the right-hand side to access the download button.

Video 1 with naïve mouse: https://osf.io/qgkva

Video 2 with CFA treated mouse: https://osf.io/wd52c

Video 3 with pSNL mouse: https://osf.io/63t4w

***Supplementary videos: Recorded Ca***^***2+***^ ***responses in L4 DRG SGCs in different conditions***

*Video examples at 8x recording speed to ease identification of Ca*^*2+*^ *responses*.

*Video 1) Naïve mouse with no activity. Video 2) CFA treated mouse 3 days after CFA injection in paw with much activity. White circle highlights an area with activity. Video 3) pSNL operated mouse 7 days after operation with some activity. White arrow points to SGC with a Ca*^*2+*^ *response. In the end of the video all SGCs have a Ca*^*2+*^ *increase caused by culling the mouse with overdose of pentobarbital*.

## References

1. Schulte A, Degenbeck J, Aue A, et al. Human dorsal root ganglia after plexus injury: either preservation or loss of the multicellular unit. BioRxiv. Published online February 8, 2023.

2. Lund H, Hunt M, Kurtovic Z, et al. A network of CD163+ macrophages monitors enhanced permeability at the blood-dorsal root ganglion barrier. BioRxiv. Published online March 27, 2023.

3. Pannese E. The structure of the perineuronal sheath of satellite glial cells (SGCs) in sensory ganglia. Neuron Glia Biol. 2010;6(1):3–10. doi:10.1017/S1740925X10000037

4. Huang TY, Cherkas PS, Rosenthal DW, Hanani M. Dye coupling among satellite glial cells in mammalian dorsal root ganglia. Brain Res. 2005;1036(1-2):42–49. doi:10.1016/j.brainres.2004.12.021

5. Vit JP, Jasmin L, Bhargava A, Ohara PT. Satellite glial cells in the trigeminal ganglion as a determinant of orofacial neuropathic pain. Neuron Glia Biol. 2006;2(4):247–257. doi:10.1017/S1740925X07000427

6. Kung LH, Gong K, Adedoyin M, et al. Evidence for Glutamate as a Neuroglial Transmitter within Sensory Ganglia. PLoS One. 2013;8(7). doi:10.1371/journal.pone.0068312

7. Hanani M, Spray DC. Emerging importance of satellite glia in nervous system function and dysfunction. Nature Reviews Neuroscience 2020 21:9. 2020;21(9):485–498. doi:10.1038/s41583-020-0333-z

8. Cherkas PS, Huang TY, Pannicke T, Tal M, Reichenbach A, Hanani M. The effects of axotomy on neurons and satellite glial cells in mouse trigeminal ganglion. Pain. 2004;110(1-2):290–298. doi:10.1016/j.pain.2004.04.007

9. Hanani M, Huang TY, Cherkas PS, Ledda M, Pannese E. Glial cell plasticity in sensory ganglia induced by nerve damage. Neuroscience. 2002;114(2):279–283. doi:10.1016/S0306-4522(02)00279-8

10. Pannese E, Ledda M, Cherkas PS, Huang TY, Hanani M. Satellite cell reactions to axon injury of sensory ganglion neurons: increase in number of gap junctions and formation of bridges connecting previously separate perineuronal sheaths. Anat Embryol (Berl). 2003;206(5):337–347. doi:10.1007/s00429-002-0301-6

11. Thakur M, Crow M, Richards N, et al. Defining the nociceptor transcriptome. Front Mol Neurosci. 2014;7. doi:10.3389/fnmol.2014.00087

12. George D, Ahrens P, Lambert S. Satellite glial cells represent a population of developmentally arrested Schwann cells. GLIA. 2018.

13. Jager SE, Pallesen LT, Lin L, et al. Comparative transcriptional analysis of satellite glial cell injury response [version 1; peer review: 1 approved]. Wellcome Open Res. 2022;7:156. doi:10.12688/wellcomeopenres.17885.1

14. Chen Z, Huang Q, Song X, et al. Purinergic signaling between neurons and satellite glial cells of mouse dorsal root ganglia modulates neuronal excitability in vivo. Pain. 2022;163(8):1636–1647. doi:10.1097/j.pain.0000000000002556

15. Avraham O, Deng PY, Jones S, et al. Satellite glial cells promote regenerative growth in sensory neurons. Nat Commun. 2020;11(1). doi:10.1038/s41467-020-18642-y

16. Mapps AA, Boehm E, Beier C, et al. Satellite glia modulate sympathetic neuron survival, activity, and autonomic function. Elife. 2022;11. doi:10.7554/eLife.74295

17. Ingram SL, Chisholm KI, Wang F, De Koninck Y, Denk F, Goodwin GL. Assessing spontaneous sensory neuron activity using in vivo calcium imaging. Published online January 17, 2023.

18. Kim YS, Anderson M, Park K, et al. Coupled Activation of Primary Sensory Neurons Contributes to Chronic Pain. Neuron. 2016;91(5):1085–1096. doi:10.1016/j.neuron.2016.07.044

19. Wang F, Bélanger E, Côté SL, et al. Sensory Afferents Use Different Coding Strategies for Heat and Cold. Cell Rep. 2018;23(7):2001–2013. doi:10.1016/j.celrep.2018.04.065

20. Chisholm KI, Khovanov N, Lopes DM, La Russa F, McMahon SB. Large Scale In Vivo Recording of Sensory Neuron Activity with GCaMP6. eNeuro. 2018;5(1):ENEURO.0417-17.2018. doi:10.1523/ENEURO.0417-17.2018

21. Zhang X, Chen Y, Wang C, Huang LYM. Neuronal somatic ATP release triggers neuron-satellite glial cell communication in dorsal root ganglia. Proc Natl Acad Sci U S A. 2007;104(23):9864–9869. doi:10.1073/pnas.0611048104

22. Gu Y, Chen Y, Zhang X, Li GW, Wang C, Huang LYM. Neuronal soma-satellite glial cell interactions in sensory ganglia and the participation of purinergic receptors. Neuron Glia Biol. 2010;6(1):53–62. doi:10.1017/S1740925X10000116

23. Suadicani SO, Cherkas PS, Zuckerman J, Smith DN, Spray DC, Hanani M. Bidirectional calcium signaling between satellite glial cells and neurons in cultured mouse trigeminal ganglia. Neuron Glia Biol. 2010;6(1):43–51. doi:10.1017/S1740925X09990408

24. Kushnir R, Cherkas PS, Hanani M. Peripheral inflammation upregulates P2X receptor expression in satellite glial cells of mouse trigeminal ganglia: A calcium imaging study. Neuropharmacology. 2011;61(4):739–746. doi:10.1016/j.neuropharm.2011.05.019

25. Blum E, Procacci P, Conte V, Hanani M. Systemic inflammation alters satellite glial cell function and structure. A possible contribution to pain. Neuroscience. 2014;274. doi:10.1016/j.neuroscience.2014.05.029

26. Ceruti S, Fumagalli M, Villa G, Verderio C, Abbracchio MP. Purinoceptor-mediated calcium signaling in primary neuron-glia trigeminal cultures. Cell Calcium. 2008;43(6). doi:10.1016/j.ceca.2007.10.003

27. Jager SE, Pallesen LT, Richner M, et al. Changes in the transcriptional fingerprint of satellite glial cells following peripheral nerve injury. Glia. 2020;68(7). doi:10.1002/glia.23785

28. Mohr, Pallesen, Richner, Vaegter. Discrepancy in the Usage of GFAP as a Marker of Satellite Glial Cell Reactivity. Biomedicines. 2021;9(8). doi:10.3390/BIOMEDICINES9081022

29. Chudler EH, Anderson LC, Byers MR. Trigeminal ganglion neuronal activity and glial fibrillary acidic protein immunoreactivity after inferior alveolar nerve crush in the adult rat. Pain. 1997;73(2):141–149. doi:S0304-3959(97)00088-2 [pii]

30. Wang F, Xiang H, Fischer G, et al. HMG-CoA synthase isoenzymes 1 and 2 localize to satellite glial cells in dorsal root ganglia and are differentially regulated by peripheral nerve injury. Brain Res. 2016;1652:62–70. doi:10.1016/j.brainres.2016.09.032

31. Takeda M, Takahashi M, Nasu M, Matsumoto S. Peripheral inflammation suppresses inward rectifying potassium currents of satellite glial cells in the trigeminal ganglia. Pain. 2011;152(9):2147–2156. doi:10.1016/j.pain.2011.05.023

32. Vit JP, Ohara PT, Bhargava A, Kelley K, Jasmin L. Silencing the Kir4.1 potassium channel subunit in satellite glial cells of the rat trigeminal ganglion results in pain-like behavior in the absence of nerve injury. J Neurosci. 2008;28(16):4161–4171. doi:10.1523/JNEUROSCI.5053-07.2008

33. Maruoka H, Kubota K, Kurokawa R, Tsuruno S, Hosoya T. Periodic organization of a major subtype of pyramidal neurons in neocortical layer V. Journal of Neuroscience. 2011;31(50). doi:10.1523/JNEUROSCI.3117-11.2011

34. Richner M, Jager SB, Siupka P, Vaegter CB. Hydraulic extrusion of the spinal cord and isolation of dorsal root ganglia in rodent. Journal of Visualized Experiments. 2017;119(e55226).

35. Seltzer Z, Dubner R, Shir Y. A novel behavioral model of neuropathic pain disorders produced in rats by partial sciatic nerve injury. Pain. 1990;43(2):205–218. doi:10.1016/0304-3959(90)91074-S

36. Goodwin G, McMurray S, Stevens EB, Denk F, McMahon SB. Examination of the contribution of Nav1.7 to axonal propagation in nociceptors. Pain. 2022;163(7). doi:10.1097/j.pain.0000000000002490

37. Pachitariu M, Stringer C, Dipoppa M, et al. Suite2p: beyond 10,000 neurons with standard two-photon microscopy. bioRxiv. Published online January 1, 2017:061507. doi:10.1101/061507

38. Stringer C, Wang T, Michaelos M, Pachitariu M. Cellpose: a generalist algorithm for cellular segmentation. Nat Methods. 2021;18(1):100–106. doi:10.1038/s41592-020-01018-x

39. Chiou J. Flexible and Fast Spike Raster Plotting. MATLAB Central File Exchange.

40. Indra AK, Warot X, Brocard J, et al. Temporally-controlled site-specific mutagenesis in the basal layer of the epidermis: Comparison of the recombinase activity of the tamoxifen-inducible Cre-ER(T) and Cre-ER(T2) recombinases. Nucleic Acids Res. 1999;27(22). doi:10.1093/nar/27.22.4324

41. Feil R, Wagner J, Metzger D, Chambon P. Regulation of Cre Recombinase Activity by Mutated Estrogen Receptor Ligand-Binding Domains. Biochem Biophys Res Commun. 1997;237(3):752–757. doi:10.1006/bbrc.1997.7124

42. Rigaud M, Gemes G, Barabas ME, et al. Species and strain differences in rodent sciatic nerve anatomy: implications for studies of neuropathic pain. Pain. 2008;136(1-2):188–201. doi:10.1016/j.pain.2008.01.016

43. Djouhri L. Spontaneous Pain, Both Neuropathic and Inflammatory, Is Related to Frequency of Spontaneous Firing in Intact C-Fiber Nociceptors. Journal of Neuroscience. 2006;26(4):1281–1292. doi:10.1523/JNEUROSCI.3388-05.2006

44. Schulte A, Lohner H, Degenbeck J, et al. Unbiased analysis of the dorsal root ganglion after peripheral nerve injury: no neuronal loss, no gliosis, but satellite glial cell plasticity. Pain. 2023;164(4):728–740. doi:10.1097/j.pain.0000000000002758

45. Chen Z, Zhang C, Song X, et al. BzATP Activates Satellite Glial Cells and Increases the Excitability of Dorsal Root Ganglia Neurons In Vivo. Cells. 2022;11(15):2280. doi:10.3390/cells11152280

46. Sekiguchi KJ, Shekhtmeyster P, Merten K, et al. Imaging large-scale cellular activity in spinal cord of freely behaving mice. Nat Commun. 2016;7(1):11450. doi:10.1038/ncomms11450

47. Spray DC, Hanani M. Gap junctions, pannexins and pain. Neurosci Lett. 2019;695:46–52. doi:10.1016/j.neulet.2017.06.035

